# Recyclable CRISPR/Cas9 mediated gene disruption and deletions in *Histoplasma*

**DOI:** 10.1101/2023.07.05.547774

**Authors:** Bastian Joehnk, Nebat Ali, Mark Voorhies, Keith Walcott, Anita Sil

## Abstract

Targeted gene disruption is challenging in the dimorphic fungal pathogen *Histoplasma* due to the low frequency of homologous recombination. Transformed DNA is either integrated ectopically into the genome or maintained extra chromosomally by *de novo* addition of telomeric sequences. Based on a system developed in *Blastomyces*, we adapted a CRISPR/Cas9 system to facilitate targeted gene disruption in *Histoplasma* with high efficiency. We express a codon-optimized version of Cas9 as well as guide RNAs from a single ectopic vector carrying a selectable marker. Once the desired mutation is verified, one can screen for isolates that have lost the Cas9 vector by simply removing the selective pressure. Multiple mutations can then be generated in the same strain by retransforming the Cas9 vector carrying different guides. We used this system to disrupt a number of target genes including *RYP2* and *SRE1* where loss-of-function mutations could be monitored visually by colony morphology or color, respectively. Interestingly, expression of two guide RNAs targeting the 5’ and 3’ ends of a gene allowed isolation of deletion mutants where the sequence between the guide RNAs was removed from the genome. Whole-genome sequencing showed that the frequency of off-target mutations associated with the Cas9 nuclease was negligible. Finally, we increased the frequency of gene disruption by using an endogenous *Histoplasma* regulatory sequence to drive guide RNA expression. These tools transform our ability to generate targeted mutations in *Histoplasma*.

**Importance:** *Histoplasma* is a primary fungal pathogen with the ability to infect otherwise healthy mammalian hosts, causing systemic and sometimes life-threatening disease. Thus far, molecular genetic manipulation of this organism has utilized RNA interference, random insertional mutagenesis, and a homologous recombination protocol that is highly variable and often inefficient. Targeted gene manipulations have been challenging due to poor rates of homologous recombination events in *Histoplasma*. Interrogation of the virulence strategies of this organism would be highly accelerated by a means of efficiently generating targeted mutations. We have developed a recyclable CRISPR/Cas9 system that can be used to introduce gene disruptions in *Histoplasma* with high efficiency, thereby allowing disruption of multiple genes.

## Introduction

The fungal order Onygenales includes multiple thermally dimorphic mammalian pathogens capable of infecting healthy hosts (1). These organisms include *Histoplasma spp.*, which are the most common cause of fungal respiratory infections in the US. Approximately 60-90% of individuals residing in the Ohio and Mississippi River Valleys are thought to have been exposed to *Histoplasma*, with the disease histoplasmosis reaching an incidence of up to 4.3 cases per 100,000 population in endemic regions and a mortality rate of up to 7% (2, 3). *Histoplasma* grows saprophytically with a filamentous morphology in the soil that generates asexual spores termed macro- and microconidia.

These conidia, the infectious agents of *Histoplasma*, are inhaled by mammalian hosts and phagocytosed by alveolar macrophages. Upon the shift to higher body temperature, *Histoplasma* converts to a pathogenic yeast form that expresses virulence factors, enabling proliferation inside the phagolysosome and eventually leading to the lysis of the infected macrophage. Thus, the dimorphic nature of this fungus is thought to be an important pathogenicity factor and is a subject of major interest (4). Several molecular tools have been developed to study morphogenesis, such as genome wide expression studies, genetic screens based on random insertional mutagenesis, and RNA interference. These techniques led to the identification of key players of dimorphic switching such as the Ryp proteins, which are required for yeast phase growth of *Histoplasma*. However, in contrast to the majority of fungal model organisms, gene disruptions or replacements in *Histoplasma* are impeded by its low rate of homologous recombination, making gene targeting challenging (5).

In recent years, CRISPR/Cas9-based genome editing has become a powerful addition to the genetic tool set for molecular research of fungi. CRISPR/Cas9-based systems have been successfully adapted to genetically manipulate yeasts such as *Saccharomyces cerevisiae*, *Pichia pastoris* and *Candida albicans* as well as multiple filamentous fungal species including *Aspergilli*, *Neurospora crassa*, and *Trichoderma reesei* (6–11). More recently, CRISPR/Cas9 tools were developed for the dimorphic fungal pathogen *Blastomyces dermatitidis* (12), a close relative of *Histoplasma*. Notably, most fungal CRISPR/Cas9 based systems rely on one of two delivery systems: i) the transformation of *in vitro* assembled Cas9 sgRNA complexes called ribo<u>n</u>ucleo<u>p</u>roteins (RNPs) or ii) the expression of both Cas9 nuclease and sgRNA within the fungal cell. The transformation of Cas9 sgRNA RNPs is generally simpler as it does not require laborious strain development and has the advantage of being essentially marker free (13). However, this method is not applicable to all species and in fact we could not optimize it for *Histoplasma* (data not shown). In contrast, expression of Cas9 and sgRNAs inside the fungal cell usually requires the genomic integration of the respective expression cassettes. The integration can result in unwanted side effects, such as the disruption of a random gene at the integration site. Furthermore, expression of sgRNAs is usually driven by RNA polymerase III promoters to generate functional non-modified sgRNAs. However, RNA pol III promoters are not well defined in fungi and there is only limited information about their expression kinetics. This problem was circumvented by Nødvig et al. by developing a special expression system for the sgRNA, which utilizes a well-defined constitutive RNA polymerase II promoter for expression instead of an RNA polymerase III promoter (14). Here the sgRNA is embedded into a larger transcript, which is flanked by self-cleaving ribozyme sequences that will give rise to the mature sgRNA.

Based on this expression system and the *Blastomyces* constructs (12), we have developed a recyclable CRISPR/Cas9 system, which makes use of an inherent feature of *Histoplasma* biology, the maintenance of episomal vectors (5). We embedded expression constructs for both Cas9 nuclease and sgRNA into a larger vector that is flanked by telomeric sequences that facilitate autonomous replication in *Histoplasma* (15). Removal of the selective pressure for the vector leads to the loss of the Cas9/sgRNA plasmid, enabling multiple rounds of CRISPR/Cas9 mediated gene targeting. We have used whole-genome sequencing to further validate that this CRISPR/Cas9 system has limited potential off-target mutations. Finally, we modified the *Blastomyces* system, introducing an endogenous glyceraldehyde-3 phosphate dehydrogenase (*GAPDH*) promoter in place of the *Aspergillus* P*_gpd_* regulatory sequence, thereby increasing the efficiency of targeting. Our CRISPR/Cas9 system can be used to introduce gene disruptions in *Histoplasma* at high efficiency with the option to mutate multiple genes.

## Results

### Assembly of an episomal CRISPR/Cas9 vector system for gene targeting in *Histoplasma*

After several unsuccessful attempts to transform in vitro assembled Cas9/sgRNA RNP complexes into *Histoplasma*, we sought to apply the CRISPR/Cas9 system originally developed by Nødvig et al. to the episomal vector system we utilize for RNAi or gene expression (11). In this case transformed DNA is carried on a linear plasmid with telomeric ends, which facilitates episomal maintenance in *Histoplasma* as long as selective pressure during growth is maintained. The vector is designed to express both a selective marker gene (either *URA5* or *hph*) and Cas9 from the bidirectional *H2AB* promoter. The sgRNA cassette is essentially the same as developed by Nødvig et al. (11) and includes the well-established *gpdA* promoter and *tef1* terminator sequences from *Aspergillus nidulans*, which drive the expression of the sgRNA precursor as a standard mRNA. The sgRNA itself is composed of a protospacer sequence and tracr RNA that are responsible for binding and activating the Cas9 nuclease as well as guiding the RNP to its destination in the genome. The sgRNA is further flanked by two autocatalytic self-cleaving ribozyme sequences, which will give rise to the mature sgRNA (Fig. 1). After the introduction of a double strand break at the protospacer mediated target site and a potential indel mutation due to error-prone non-homologous end joining repair, one can screen for isolates that have lost the vector when selective pressure is removed from the media.

**Figure 1.**
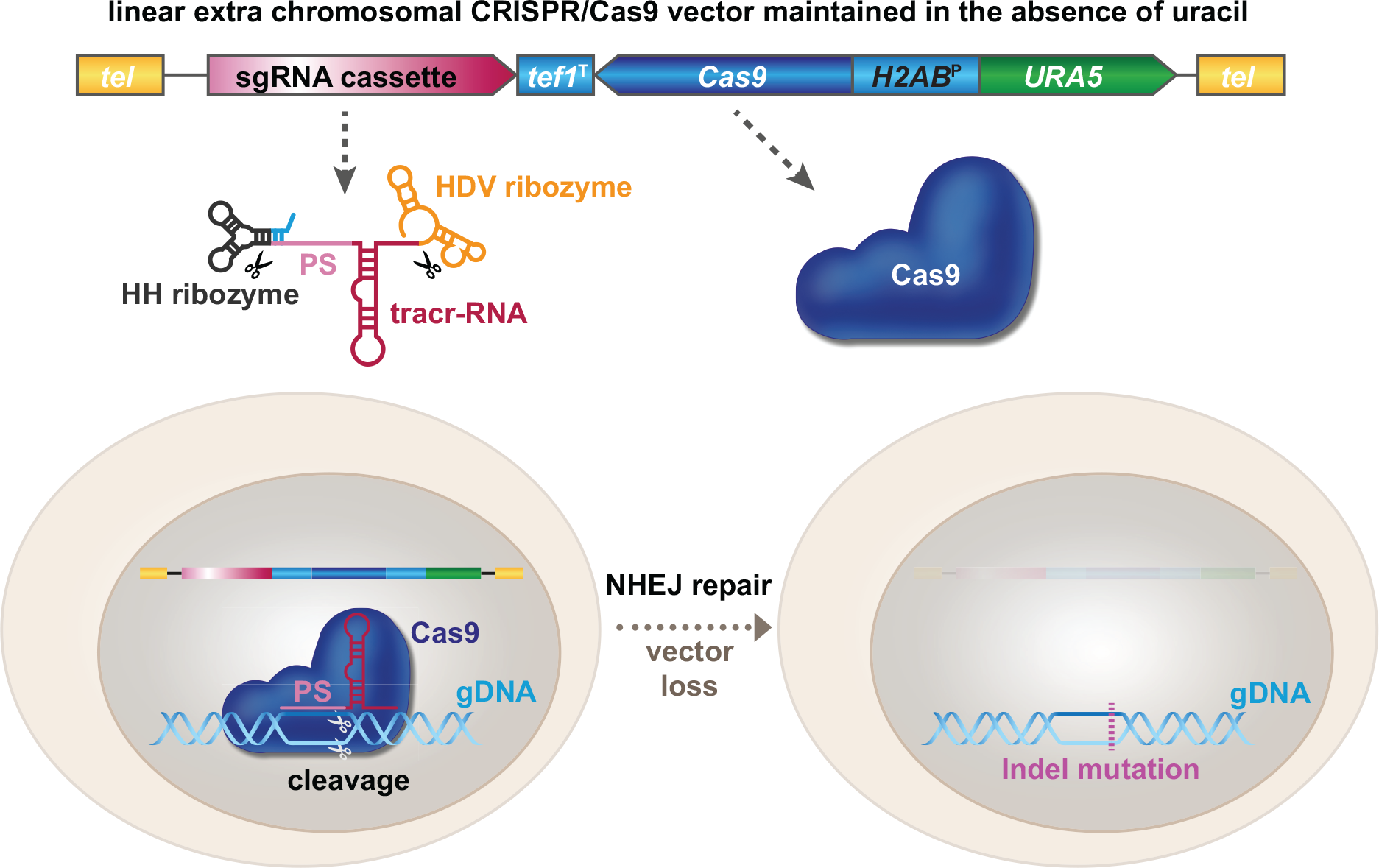
Schematic representation of the CRISPR/Cas9 mediated targeted gene disruption in *H. capsulatum*. Under selective pressure the CRISPR/Cas9 plasmid is maintained in *Histoplasma* as an extrachromosomal vector, which carries both a codon optimized version of Cas9 and the sgRNA (based on the concept of Nødvig et al., 2015). The sgRNA is expressed as a precursor driven by a constitutive RNA polymerase II promoter (*gpdA*^P^), resulting in a larger mRNA-like transcript with 5’-cap and poly-(A) tail. The mature sgRNA gets excised from its larger transcript via the action of two flanking ribozyme sequences (hammerhead, HH and hepatitis delta virus, HDV). The sgRNA then recruits the Cas9 nuclease to a protospacer (PS)-mediated target site in the genome, where it cuts the gDNA. The resulting double-strand break is repaired by the non-homologous end joining (NHEJ) mechanism, an error-prone process that often introduces small indel mutations. Growing the resultant mutant strain under non-selective conditions (*i.e.* in the presence of uracil) allows loss of the CRISPR/Cas9 vector, which then permits re-use of the *URA5* marker for subsequent transformations.

Potential protospacer sequences for target genes were identified with CRISPOR (http://crispor.tefor.net/) (16). The main criteria for protospacer selection for a specific target gene were a low off-target score as well as a location within the first exon or the first 100 bp of the coding sequence to avoid generating a mutant allele that encodes a partially functional truncated protein due to late frame shift mutations. Protospacer sequences were introduced into the sgRNA cassette via fusion PCR with overlapping primer pairs that contain the 20 bp protospacer sequence as well as a 6bp inverted repeat of the protospacer sequence to ensure correct folding/cleavage of the <u>h</u>ammer<u>h</u>ead (HH) ribozyme at the 5’-end. For final CRISPR/Cas9 vector assembly, we utilized the Gateway cloning system. We constructed two different Cas9 containing expression vectors with either *URA5* or *hph* as selection markers, which both incorporated the *ccdB* containing Gateway cassette instead of the sgRNA cassette. Newly assembled sgRNA cassettes were generated in pDONR vectors, which were then recombined with the Cas9 expression vectors. This method allows the generation of a sgRNA cassette that can easily be transferred into either CRISPR/Cas9 vector depending on the selection marker needed.

### Highly efficient gene disruption of *RYP2*

We first tried to use this system to disrupt *RYP2*, a key transcriptional regulator required for yeast phase growth at 37°C (Fig. 2A) (17). Successful disruptions of *RYP2* should be identifiable by filamentous colony morphology even under yeast promoting growth conditions. Upon electroporation of the linearized *RYP2* targeting CRISPR/Cas9 vector, 97% of the primary transformants showed at least a partial filamentous phenotype whereas control transformants carrying a Cas9 expressing vector without a sgRNA appeared exclusively in the yeast morphology (Fig. 2B). From the primary transformants we isolated a pure filamentous colony by passaging the primary colony for two generations. Both primary transformants as well as first generation passages appeared as mixed yeast-filamentous colonies, indicating mosaic colonies consisting of wild-type (yeast-form) and *ryp2* mutant (filamentous-form) cells (Fig. 2C). We validated the disruption of the *RYP2* target site in the filamentous mutant by PCR amplification of a 500bp region surrounding the expected cut site followed by Sanger sequencing. We could observe a two-base pair insertion exactly at the protospacer-mediated cut site three bp upstream of the protospacer adjacent motif (PAM) (Fig. 2D). As predicted, sequencing of the same locus from Cas9 expressing control strains without sgRNA all resulted in wild-type sequences.

**Figure 2.**
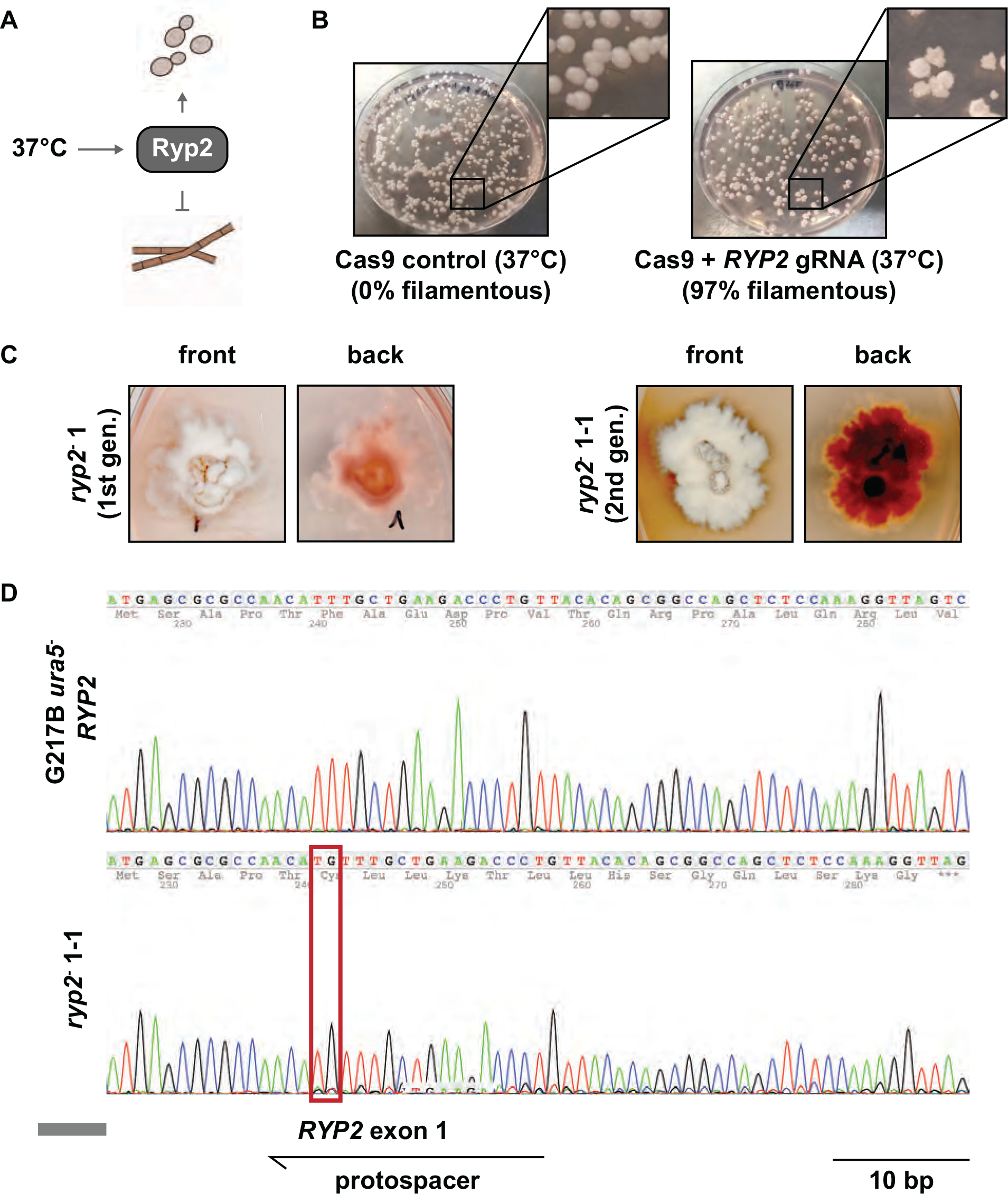
CRISPR/Cas9 mediated disruption of *RYP2*. (A) Schematic representation of *RYP2*-mediated regulation of the dimorphic switch in *H. capsulatum*. At mammalian body temperature, Ryp2 is required to promote yeast-phase growth. (B) Primary transformants carrying the CRISPR/Cas9 vector targeting *RYP2*. Whereas transformants carrying a Cas9-only expressing control vector exhibited a smooth yeast-colony shape, transformants carrying the *RYP2* targeting CRISPR/Cas9 vector showed predominantly a wrinkled/filamentous phenotype. (C) Phenotypes of first-generation and second-generation *ryp2^-^* mutants displaying the gradual increase from a mixed yeast/filamentous colony shape to a completely filamentous phenotype. Colonies also showed increased red pigmentation indicating altered secondary metabolism associated with filamentous growth. (D) Sanger sequencing of the *RYP2* target region of the WT and the second-generation *ryp2^-^* mutant revealed an insertion of TG in the protospacer-mediated target region in the mutant causing a frame-shift mutation in the first exon.

### Successive CRISPR/Cas9 disruption of multiple genes in *Histoplasma*

We investigated whether this system could be used to disrupt multiple genes in *Histoplasma* via vector recycling. We first disrupted *SRE1*, which encodes a repressor of siderophore biosynthesis. Siderophores are small high affinity iron scavengers that serve to transport iron across membranes in bacteria and fungi. When complexed with iron, siderophores confer an orange tinge to colonies, especially when siderophores are in excess as previously observed by targeting *SRE1* with RNAi (18). We reasoned that CRISPR-Cas9-generated *sre1* mutants should be easy to identify on a plate due to increased siderophore formation and thus characteristic orange pigmentation. However, all primary transformants were indistinguishable from the wild-type or Cas9-expressing controls and appeared as creamy white colonies. Similar to primary *ryp2^-^* mutant transformants, which appeared as mixed colonies of yeast and filamentous phenotype, primary transformants from the *SRE1* targeting CRISPR/Cas9 vector showed a mixed population of cells with either wild-type or the disrupted allele. To identify the frequency of successfully disrupted mutant cells we applied a method commonly used for CRISPR applications in cell lines of higher eukaryotes called <u>T</u>racking of <u>I</u>ndels by <u>De</u>composition (TIDE). TIDE identifies the frequency and efficiency of Cas9-mediated frame shift mutations in a mixed pool of CRISPR/Cas9 mediated mutants from Sanger sequencing traces (19). To identify possible frame shift mutations in the *SRE1* target region, we used colony PCR to amplify a 519 bp fragment centered around the putative Cas9 cutting site from colonies of either wild-type, Cas9-expressing controls, or mixed colonies of transformants with the *SRE1*-targeting Cas9 vector. The resulting fragments were sequenced via standard Sanger sequencing and sequence traces were uploaded to the TIDE online platform (http://tide.nki.nl/). The TIDE algorithm aligns the wild-type sequence with sequence traces of potential mutant/mixed colonies and determines the frequency and alterations in the alignment following the cutting site. The number of aberrant sequences in the mixed pool compared to the wild-type gets interpreted as efficiency of the respective sgRNA to generate mutant cells. Interestingly, our initial colonies of *sre1^-^* mutants had relatively low disruption efficiency, between 0.5% - 8.6% (Fig. 3A). Passaging the mutants with the highest disruption efficiency successively increased the efficiency of following generations until we obtained a mutant with 96.6% efficiency in generation three, thus approaching homogeneity. The TIDE algorithm further allows the identification of the CRISPR-mediated frame shift mutation, which was an insertion of a guanine at the protospacer mediated cutting site (Fig. 3A, right panel). This insertion was further confirmed via gDNA extraction and sequencing of the respective region (Fig. 3B). The resulting mutant colony was then passaged on non-selective media for three generations and individual colonies were replica-plated on selective and non-selective media to identify mutants that had lost the *SRE1* targeting Cas9 vector. Two individual colonies that carried the guanine insertion mutation and had lost the Cas9 vector were selected for further CRISPR/Cas9 experiments.

**Figure 3.**
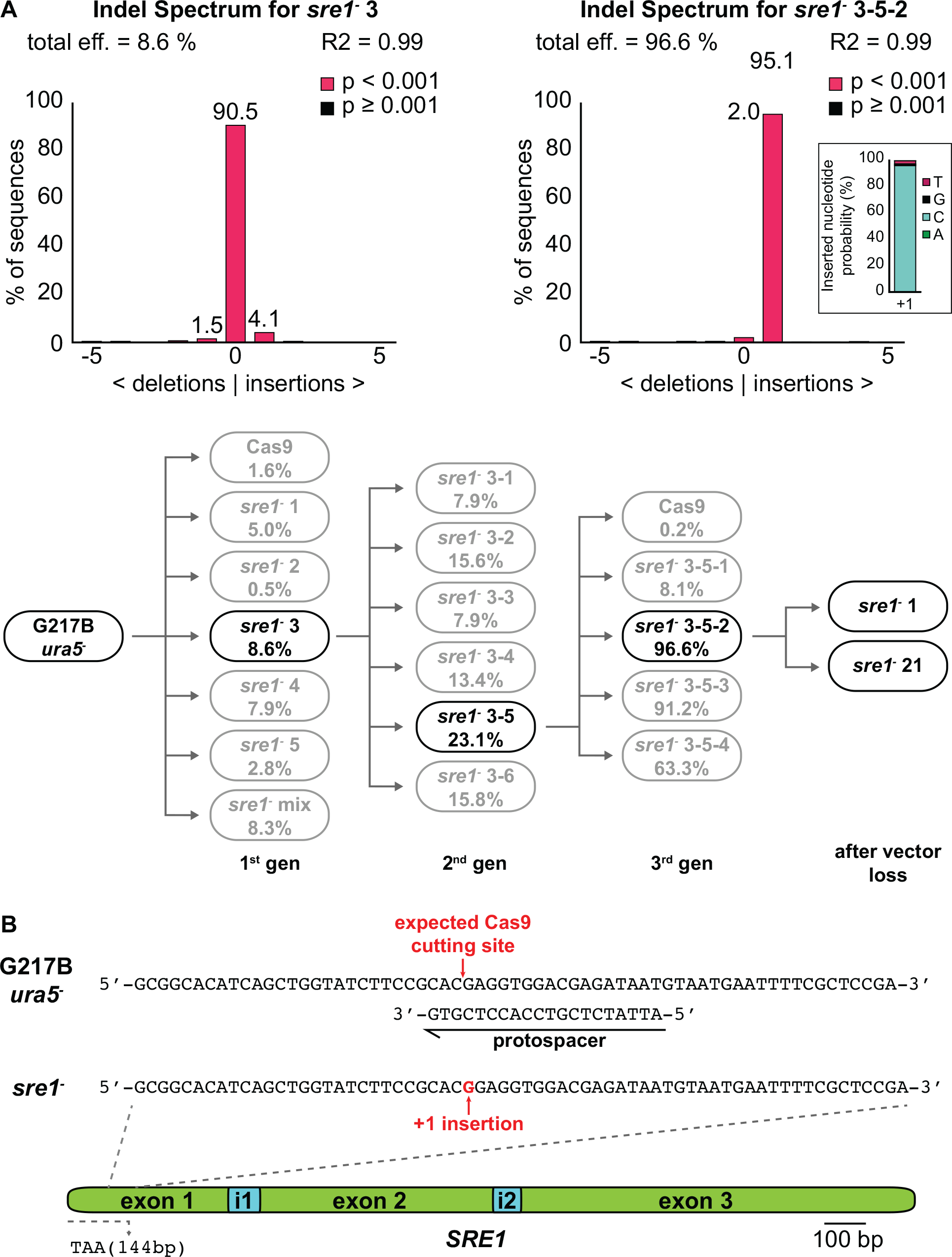
Evaluation of *sre1^-^*disruption mutants with the online platform TIDE. Transformants with a *SRE1*-targeting CRISPR/Cas9 vector either directly after transformation or after each passaging were analyzed with TIDE (https://tide.deskgen.com), which detects the predominant mutations (i.e. deletions or insertions) as well as their frequencies. To do so, approximately 500 bp of the sgRNA mediated target sequence were amplified via colony PCR from a control strain as well as mutant colonies harboring the mutation of interest. The DNA fragments were sequenced by standard Sanger sequencing and compared with the TIDE platform. This online tool aligns the first part of the sequences before the anticipated cutting site and calculates the frequency of aberrant sequences following the cutting site for the mutant colony, in this case a disruption mutant of *SRE1* (96.6%) compared to the control sequence. The bottom of panel A shows the gene-editing efficiency of the mutant isolates after subsequent generations of passaging based on TIDE analysis. The gene editing efficiency increases for most colonies after each passaging, resulting in homogeneous mutant isolates after three passages. An additional passage on non-selective medium (*i.e.* in the presence of uracil) resulted in the loss of the vector. (B) Sequencing of the final mutants after Cas9 vector loss confirmed a G insertion in the first exon of the *sre1^-^* mutants resulting in a premature stop codon after 144 bp.

As mentioned above, we expected pure *sre1^-^* mutants to display increased siderophore production, resulting in orange-pigmented colonies especially on high-iron containing media. Phenotypic characterization revealed that the mutants indeed showed increased orange pigmentation compared to wild-type (Fig. 4A). To explore the possibility of making double mutants using CRISPR, we turned to the siderophore biosynthesis pathway. The first enzyme in this pathway, L-ornithine monooxygenase, is encoded by *SID1*, which is essential for both extra- and intracellular siderophores. Sre1 normally represses *SID1* expression under iron-replete conditions (Fig. 4B) (20, 21). To verify that our CRISPR/Cas9 system can make multiple gene disruptions in the same strain, we disrupted *SID1* in the *sre1^-^* mutants followed by Cas9-vector recycling in non-selective media. The resulting *sre1^-^ sid1^-^* double mutants were confirmed by Sanger sequencing and compared with wild-type as well as *sre1^-^* and *sid1^-^*single mutants for their capacity to produce siderophores. The disruption of *SID1* in the *sre1^-^* background led to a complete loss of pigmentation (Fig. 4A), consistent with the inability of the mutant to produce siderophores (Fig. 4C).

**Figure 4.**
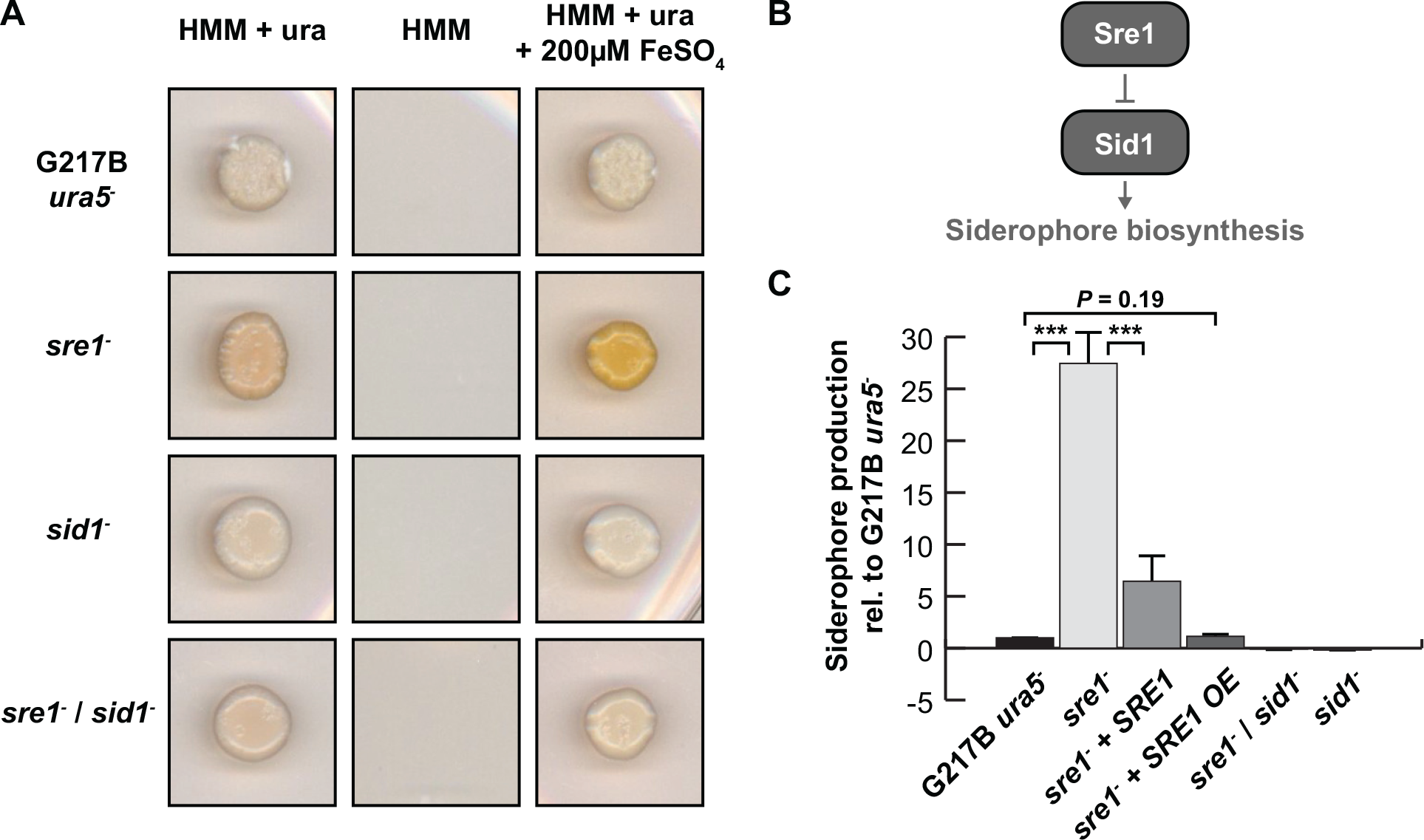
CRISPR/Cas9 vector recycling allows successive gene disruptions and complementation. (A) CRISPR/Cas9 vector recycling in the *sre1^-^* mutants allowed re-use of the *URA5* marker to target a second gene in the *sre1^-^*background. The loss of *SRE1* resulted in derepressed siderophore expression, which led to orange pigmentation of the colonies due to increased iron-bound siderophores, especially on high iron containing media. Disruption of *SID1* in the *sre1^-^* background, which encodes the first enzyme in the siderophore biosynthesis pathway (as diagrammed in B) resulted in complete abolishment of siderophore production, indicated by the loss of orange pigmentation of the colonies of the *sre1^-^ sid1^-^* double mutants. (C) After loss of the Cas9 plasmid in the *sre1^-^* mutant, the *URA5* marker gene was available for selection of complementation plasmids. Siderophore production was analyzed with the CAS assay and showed partial complementation when *SRE1* was expressed under its native promoter and almost complete complementation when *SRE1* was overexpressed under the control of the constitutive *GAPDH* promoter (OE). In contrast, both *sid1^-^* and *sre1^-^ sid1^-^* double mutants showed no siderophore production as expected.

The selection marker that was originally used to select for the Cas9 vector to generate the *sre1*^-^ mutant was re-purposed to complement the mutant. We introduced a genomic copy of the *SRE1* wild-type locus as well as an overexpression construct in which the expression of *SRE1* was driven by the constitutive *GAPDH* promoter into the *sre1^-^* mutant and analyzed siderophore production levels by <u>c</u>hrome <u>a</u>zurol <u>S</u> (CAS) assay (Fig. 4C). We observed partial complementation of repression of siderophore biosynthesis when *SRE1* was expressed under its native promoter and complete complementation when it was constitutively expressed under the control of the *GAPDH* promoter. As expected, both *sid1^-^* and *sre1^-^ sid1^-^* double mutants were unable to produce siderophores (Fig. 4C).

### Whole genome sequencing revealed slightly increased SNP accumulations in Cas9 expressing strains

To fully characterize the genotypes of the first round of CRISPR strains, we subjected them to deep sequencing along with several control strains derived from the same frozen WU15 stock as the *sre1^-^* CRISPR strains. These isolates included strains transformed with a Cas9 control vector (Cas9-1 and Cas9-2), a strain passaged without transformation (G217B_ura5_old), and a strain grown in liquid culture for genomic DNA purification without passaging (G217B_ura5_new). The relationships among the sequenced strains are illustrated in Fig. 5.

**Figure 5.**
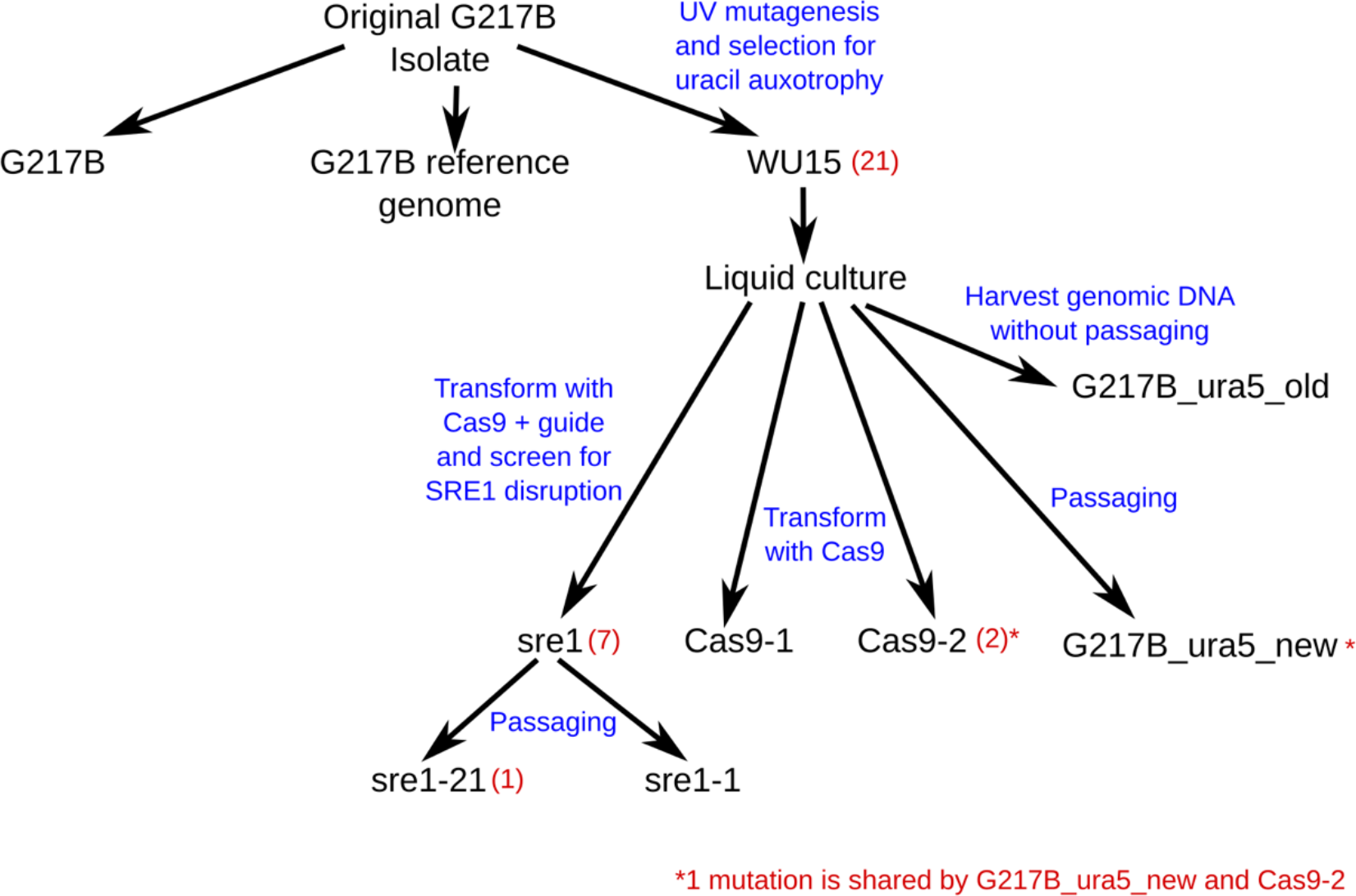
Validation of the CRISPR/Cas9 system via whole genome sequencing. We compared the genomes of two CRISPR/Cas9 generated mutants (*sre1-1*, *sre1-21*) with Cas9 only expressing control strains (*Cas9-1*, *Cas9-2*) and the parental strain from which we extracted gDNA for deep sequencing either before (G217B_ura5_old) or after passaging along with the CRISPR mutants (G217B_ura5_new). The diagram shows the strain lineages and whether strains were transformed and passaged for gene-disruption or control. Strain unique variant are given in red parenthesis including the targeted CRISPR/Cas9 mutation.

Based on comparison among strains, we discarded 917 variant positions that could not be distinguished among our sequencing samples, including 314 positions representing clear errors in the reference. This left 36 variant positions, including the targeted CRISPR mutation in *SRE1* (shared by *sre1-1* and *sre1-21*). Strain-unique variant positions, corresponding to mutations acquired after limited passaging, are quite rare: 1 in *sre1-21* and 2 in Cas9-1. An additional 6 variants (2 insertions, 2 adjacent polyG deletions, and 2 point mutations) are shared by sre1-1 and sre1-21. There are 21 variants common to the WU15-derived strains, of which 18 are adjacent to a 3617 bp deletion excising 3 genes including *URA5* (Fig. S1).We attribute both the point mutations and the large deletion to the UV mutagenesis that produced the WU15 strain. Although this strain has been previously reported and is widely used, this is the first precise characterization of the genomic change responsible for the Ura^-^ phenotype. Finally, there is one short sequence duplication shared by G217B_ura5_new and Cas9-2.

We observed no large deletions in non-repeat regions for Cas9 transformed strains, with or without CRISPR guides.

### Cas9 dual sgRNA vectors facilitate complete gene deletions

The method described thus far generates gene disruptions rather than deletions. However, during the course of protocol optimization, we noticed that introducing two sgRNAs into the CRISPR/Cas9 vector that target the same locus resulted in a deletion of the fragment between the protospacer-mediated cut sites. We decided to explore this phenomenon further by using two sgRNAs with protospacer sequences targeting the start and end of a whole coding region to facilitate complete gene deletions. We designed two sgRNA cassettes that target the 5’- and the 3’-end of the coding sequence for the velvet protein Vea1. *VEA1* RNAi studies have shown that Vea1 is required for appropriate dimorphic switching upon temperature change (22). Primary transformants were screened by PCR using a primer that would anneal to the *VEA1* 5’- and 3’-flanking regions, resulting in two different bands based on the absence or presence of the *VEA1* CDS (Fig. 6A). Selected colonies were passaged and screened by PCR until the *VEA1* wild-type band disappeared. The deletion of the *VEA1* CDS in the mutants was validated by Southern hybridization using probes in either the 5’UTR flanking region of *VEA1* (Fig. 6B) or the *VEA1* CDS (Fig. S2). Phenotypical characterization of the *vealΔ* mutants confirmed the results of Laskowski-Peak et al. (22), which showed a faster rate of filamentation in *VEA1* RNAi strains at room temperature compared to wild-type. Similarly, we observed the appearance of white, filamentous patches much earlier for the *vea1Δ* strains compared to the parental strain (Fig. 6C).

**Figure 6.**
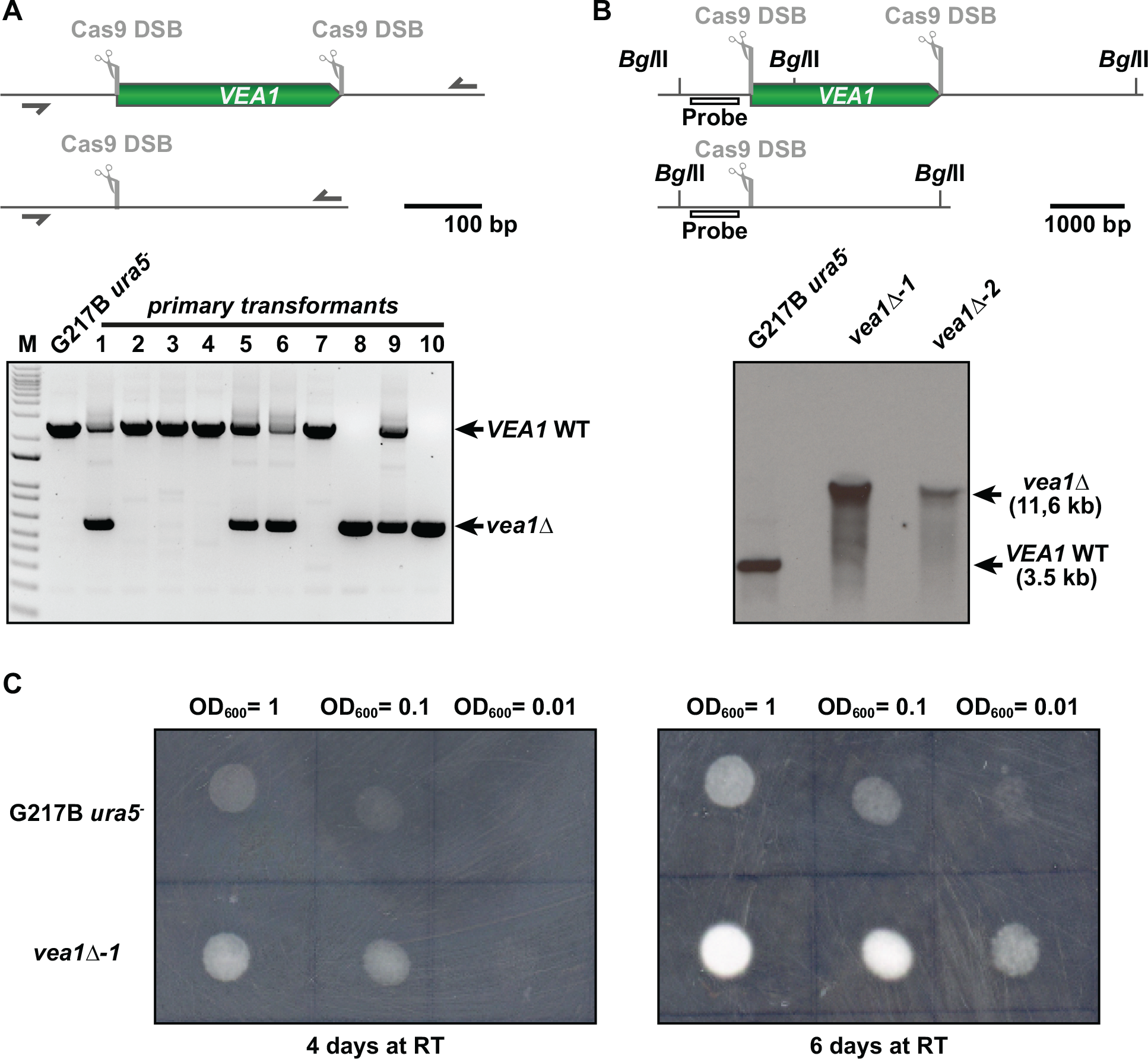
sgRNA multiplexing for complete gene deletions. We introduced two sgRNA cassettes into the Cas9 expressing vector, targeting the 5’- and 3’-end of the *VEA1* CDS respectively. Colony PCR of primary transformants (lower part of panel A) resulted in two bands for 4 of 10 isolates and only one band for two isolates, indicating partial or complete loss of the *VEA1* CDS in those mutants. WT indicates wild-type. (B) Southern hybridization of *vea1*Δ mutants confirmed the deletion of the *VEA1* CDS in two independent mutants. Location of the probe is shown in the upper part of panel B and the Southern blot is shown in the lower part of panel B. WT indicates wild-type. (C) Phenotypical characterization of the mutants grown at room temperature (RT) was performed by spotting dilutions of wild-type (G217B *ura5^-^*) and *vea1Δ* mutant strains on plates. The *vea1*Δ mutants displayed accelerated filamentation as compared to the parental strain as indicated by the increased density of fluffy white filamentous growth. The difference between wild-type and mutant strains was most pronounced after six days of incubation.

### Modulating sgRNA expression alters the frequency of Cas9-mediated editing

To further optimize Cas9-mediated gene editing, we assessed the effect of expressing sgRNAs under the control of a native *Histoplasma* promoter. We constructed episomal Cas9 plasmids driving the expression of an identical mCherry sgRNA under the control of either the *A. nidulans GAPDH* promoter (P_gpdA_) adapted from Nødvig *et al.* 2015 or the native *Histoplasma* glyceraldehyde 3-phosphate dehydrogenase (*GAPDH*) promoter (P_GAPDH_) (Fig. 7A). These plasmids were then transformed into a *Histoplasma* strain (mCherry HcG217B (23)) harboring an integrated mCherry marker driven under the expression of the *CBP1* promoter. The robust mCherry expression in this strain results in yeast colonies that are bright pink by eye with levels of fluorescence that are detectable via microscopy and flow cytometry (Fig. 7B). Following transformation, we initially observed that a majority of P_GAPDH_ transformants were cream colored by eye in contrast to P_gpdA_ transformants that retained some degree of pink color. Following these preliminary observations, two P_GAPDH_ isolates and two P_gpdA_ isolates were selected at random and passaged under selection in liquid culture. These isolates were subsequently used for Sanger sequencing and flow cytometry to quantify editing efficiency and mCherry signal, respectively. Interestingly, our TIDE analysis revealed that in contrast to the average 25% editing efficiency observed in P_gpdA_ transformants, P_GAPDH_ transformants displayed an average editing efficiency of 93% (Fig. 7C). This finding was also corroborated by our flow cytometry data, where 100% of P_GAPDH__1 and 94.1% of P_GAPDH__2 cells were negative for mCherry signal (Fig. 7D). These findings suggest that optimizing expression of sgRNAs through use of a native promoter can enhance editing efficiency and minimize the passaging time required to generate a pure mutant.

**Figure 7.**
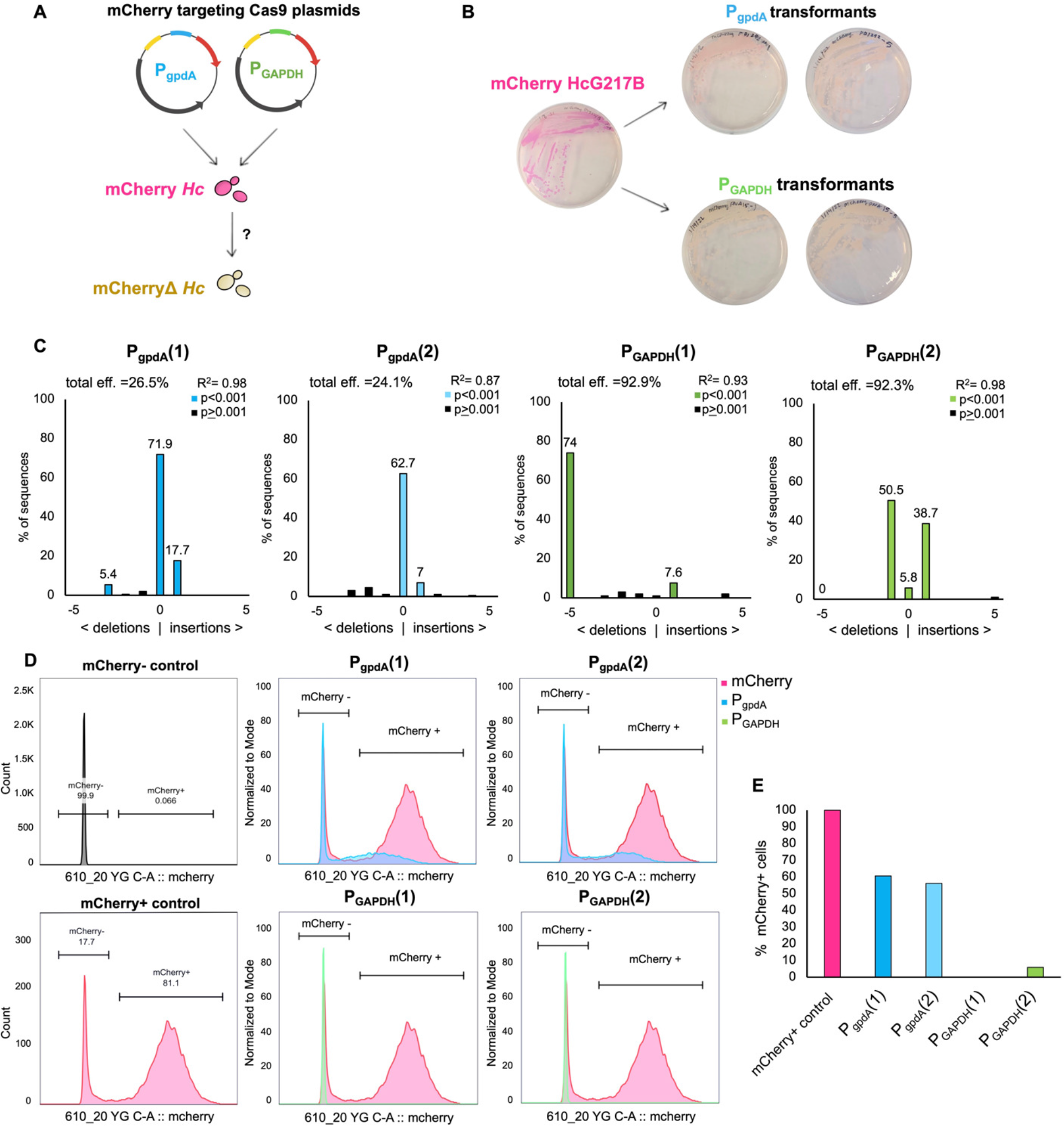
Cas9-sgRNAs expressed under the control of the *Histoplasma* GAPDH promoter showed increased editing efficiency. A) Schematic displaying experimental approach. Plasmids containing either the *A. nidulans* GAPDH promoter adapted from Nødvig et al. 2015 (P_gpdA_) or the native *Histoplasma* GAPDH promoter (P_GAPDH_) driving the expression of an identical mCherry targeting guide RNA were transformed into the mCherryHcG217B strain and used to assess differences in editing efficiencies. B) Plate images of the parental mCherryHcG217B strain alongside transformants selected from each plasmid transformation that were subsequently used for Sanger sequencing and flow cytometry. C) Indel spectrum of two individual isolates obtained from each plasmid transformation. Sanger sequencing of the mCherry locus was used to conduct TIDE analysis (https://tide.nki.nl/) to quantify the frequency of insertions and deletions following the predicted cut site. D) Histograms displaying measurements of mCherry signal acquired through flow cytometry. Overlay plots display normalized counts for each isolate in comparison to the starting mCherryHcG217B strain. E) Quantification of flow data displaying percentage of mCherry+ cells within each isolate.

## Discussion

Here we describe the development of a recyclable CRISPR/Cas9 system which can be used to introduce gene disruptions or deletions in *Histoplasma*, without leaving major marks in the genome other than the targeted mutations. As proof of principle, we generated disruptions of *RYP2*, *SRE1*, and *SID1*, as well as a complete deletion of *VEA1*. This method allows the rapid generation of targeted mutants with a very low risk for off-target mutations. Furthermore, multiplexing sgRNA expression cassettes in the Cas9 expression vector can be used to successfully delete whole genes and has the potential to accelerate gene disruptions for multiple loci.

The molecular toolbox for studying the basic biology of *Histoplasma* includes genome wide expression studies, genetic screens based on random insertional mutagenesis, and RNA interference. However, in contrast to other fungi, targeted gene disruptions or replacements are rare, due to the low rate of homologous recombination (24). In the present study we built on technology applied in *Blastomyces* (12) to demonstrate that CRISPR/Cas9 technology can be utilized to generate targeted gene disruptions and deletions in *Histoplasma*. We utilized an inherent feature of *Histoplasma*, the maintenance of extra chromosomal DNA, and embedded both Cas9 and sgRNA expression cassettes into an episomal vector system that can be recycled after successful gene disruption. Initial genome editing efficiency after transformation of the Cas9- sgRNA constructs was not always high in *Histoplasma*, resulting in colonies with mixed populations of wild-type and mutated cells, but successive passaging led to rapid increase in editing efficiency resulting in homogenous mutated colonies after 2-4 passages. This is in contrast to the CRISPR/Cas9 strategy applied in the dimorphic pathogen *Blastomyces dermatitidis* where Kujoth et al. could not detect increased genome editing upon successive passaging (12). Interestingly, expressing Cas9-sgRNA constructs under the control of the *Histoplasma GAPDH* promoter rather than the *A. nidulans GAPDH* promoter led to increased efficiency of targeting in *Histoplasma*.

We found that the combination of colony PCR and the TIDE algorithm simplifies and streamlines the screening process of primary transformants and passaged isolates to obtain targeted disruptions. It was most helpful to use TIDE as a tool to select transformants with highest editing efficiency for further passaging to isolate homogeneous mutant colonies. Additionally, we have used TIDE to evaluate general efficiency of different sgRNAs targeting the same genomic locus. This screening is particularly helpful because the efficiency of individual sgRNAs can vary widely for unknown reasons. The G/C content of the protospacer as well as the four bases preceding the PAM sequence seem to have a huge influence on the efficiency of sgRNAs (25, 26). Choosing the most desirable sgRNA can become a compromise between location of the protospacer sequence in the target region and the predicted efficiency of the sgRNA based on its sequence. TIDE can be used after transformation to identify the most efficient sgRNA for a locus with limited protospacer availability.

The delivery of Cas9-sgRNA RNPs into fungal cells is usually achieved either via transformation of *in vitro* assembled RNPs or by integration of expression cassettes for Cas9 and sgRNA into the genome (9, 12, 13, 27, 28). Concerns about potential off-target effects due to irregular Cas9 activity have led to the development of transient Cas9 expression systems, where Cas9 and sgRNA expression cassettes can be removed from the genome after successful gene editing (10, 11, 29). A transient expression system bears several advantages, the first being the ability to reuse selectable marker genes, especially in fungi with limited marker availability. In *Histoplasma*, there are currently three marker genes with documented use: *URA5* (orotate phosphoribosyltransferase), *hph* (hygromycin phosphotranseferase), and *Sh ble* (bleomycin/Zeocin resistance gene). Of these, *URA5* requires the auxotrophic *ura5^-^*background strain, leaving just two marker genes for genetic manipulations in the wild-type strain. Another advantage of a recyclable transient expression system is the elucidation of complex genetic networks by consecutive gene disruptions or deletions as exemplified here by the successive deletion of *SRE1* and *SID1*. Additionally, loss of the Cas9-sgRNA expression construct allows introduction of a complementation clone without concern that the complementation sequence will be targeted by Cas9. Recycling of the Cas9 nuclease further prevents the risk of potential off-target mutations, which could accumulate due to prolonged Cas9 expression. Interestingly, potential cytotoxic effects due to Cas9 expression in fungi are a matter of debate. Whereas it seems that some fungal groups of the genera *Aspergillus, Candida, and Cryptococcus* are not affected by prolonged Cas9 expression, cytotoxic effects have been shown in *Saccharomyces* and *Candida glabrata* (30). However, it is still unclear whether it is the Cas9 expression itself that actually results in increased off-target mutations. We investigated this by comparing the genomes of CRISPR/Cas9 generated mutants to Cas9-only expressing strains and wild-type strains. We observed a slight increase in variants in the genome of the CRISPR-generated mutants compared to wild-type. Five of these variants were strain-unique and 6 were shared among the CRISPR edited mutants. Further experimentation will determine whether these latter 6 mutations will continue to be overrepresented in strains expressing CRISPR/Cas9. Another concern that has been raised for CRISPR-based applications in cells of higher eukaryotes is the potential for large-scale genomic rearrangements in the target region (31). However, we did not observe any large deletions or rearrangements in any of the sequenced isolates. Our results confirm previous observations that CRISPR/Cas9 applications in fungi are generally a robust and suitable alternative to classical molecular biology methods, such as homologous recombination. The approach described here is likely to have a transformative effect on dissecting *Histoplasma* biology.

## Materials and Methods

### *Histoplasma* strains and culture conditions

All experiments were carried out in the *Histoplasma* G217B *ura5^-^* background. A list of generated strains can be found in Table S3. Yeast cultures of *Histoplasma* strains were propagated in liquid <u>H</u>istoplasma <u>m</u>acrophage <u>m</u>edium (HMM) (32) supplemented with uracil (200 µg/ml), hygromycin (200 µg/ml) or Zeocin (50 µg/ml) as indicated. Liquid cultures were grown at 37°C with 5% CO_2_ on an orbital shaker with 120 rpm. For phenotypical characterization of the strains, yeast cultures were diluted to an OD_600_ = 1 and 10 µl were spotted on solid HMM plates, which were incubated at 37°C with 5% CO_2_.

For cloning purposes either *E. coli* DH5α or One Shot® *ccdB* Survival^TM^ 2 T1^R^ (Invitrogen) for plasmids containing the *ccdB* containing Gateway cassette were used.

### Generation of episomal CRISPR/Cas9 vectors

All plasmids and primers used in this study are described in Table S1 and S2. Based on the work of Nødvig et al. (11), we created an episomal vector that expresses fungal codon optimized *cas9* from *Streptococcus pyogenes* as well as a hybrid sgRNA. For the vector backbone we used pBJ209, which was amplified from the RNAi vector pSB23 with primers OAS5744/OAS5745, which introduce ApaI and NheI restriction sites respectively. Next, we amplified the *ccdB* containing Gateway cassette with primers OAS5734/OAS5735 from pSB23, which introduced ClaI restriction sites at 5’- and 3’-ends and cloned the Gateway cassette into the ClaI site of pBJ209, resulting in pBJ213. The codon optimized *cas9* sequence was amplified from pPTS608-Cas9-hyg (12) with primers OAS5736/OAS5737 that introduced NheI- and ApaI-sites at the 5’- and 3’-end as well as attB-sites for subcloning in pDONR. The *cas9* sequence was excised from its larger amplicon with NheI and ApaI and cloned into NheI/ApaI digested pBJ213 to make pBJ219. Any gRNA cassette cloned into the pDONR vector can be recombined with pBJ219 via Gateway cloning to produce a final CRISPR/Cas9 targeting vector for transformation into *Histoplasma*. Details for individual targeting vectors are given in supplemental material.

### Evaluation of protospacer sequences

Protospacer sequences specific for the Sp-Cas9 (20bp-NGG) were designed to target the first exon of the target genes to disrupt the coding sequence as far upstream as possible. Potential candidates were evaluated for specificity and possible off-target sites with the online tool CRISPOR (http://crispor.tefor.net) (16).

### TIDE analysis of heterogeneous colonies

To determine the editing efficiency of the respective gRNAs, we used the online tool TIDE (http://tide.nki.nl/) (19). Colonies of positive transformants as well as wild-type controls were subjected to colony PCR to amplify an approximately 500bp fragment of the gRNA mediated target region centered around the putative cutting site. For colony PCR, *Histoplasma* yeast colonies were lysed by boiling in 100 µl 0.02 M NaOH for 10 min. 2 µl of the supernatant was directly used as template for standard PCR reactions with either Phusion or Taq polymerase. The fragments were sequenced by Sanger DNA sequencing. The sequence traces of wild-type and heterogeneous mutated colonies were aligned with the TIDE online tool, which identifies mutations introduced at or near the putative cutting site and determines their approximate frequency in a heterogeneous cell population.

### CAS assay for siderophore quantification

The siderophore production level of different mutants was determined using <u>c</u>hrome <u>a</u>zurol <u>S</u> (CAS) as previously described (21). *Histoplasma* yeast cultures, grown in triplicate in liquid HMM, were pelleted, washed with PBS and resuspended in RPMI without phenol red to avoid potential interference with the colorimetric assay. The cultures were grown for 24 h at 37°C. The culture supernatant was mixed 1:1 with modified CAS assay solution (0.6 M HDTMA, 15 µM FeCl_3_, 150 µM CAS, 0.5 M MES, 3.4 mM 5-sulfosalicyclic acid) (33) in a 96 well plate and incubated for 3-4 h. OD_630_ was measured with a plate reader and normalized to OD_600_ of the cultures to account for cell density.

### Whole genome library preparation and sequencing

Dual-indexed paired-end libraries for whole genome sequencing were prepared with the Nextera DNA Flex Library Prep Kit from Illumina. Individual libraries were barcoded with Nextera DNA CD Indexes (Illumina) and analyzed for average size distribution and concentration on a High Sensitivity DNA Bioanalyzer chip from Agilent Technologies. Fragments of individual libraries had an average size distribution of 500-550 bp and 5 ng of each library were pooled for multiplexed sequencing. All 6 strains were sequenced in a single lane on an Illumina HiSeq 4000 at the UCSF Center for Advanced Technology (UCSF CAT) yielding 101-mer paired-end reads with ∼400x coverage of the genome per strain. To distinguish mutations in the WU15 parental strain from errors in the reference, we additionally analyzed reads from wild-type G217B (34) with ∼200x coverage of the genome.

### NGS data analysis

The raw data are available at the NCBI Sequence Read Archive (SRA) under SRA accession PRJNA971667. The reads were analyzed as illustrated in Supplemental Fig. S3. In summary, reads were aligned to the 11/30/2004 version of the G217B genome assembly from the Genome Sequencing Center (GSC) at Washington University as mirrored at https://histo.ucsf.edu/downloads/ using BWA-MEM (35) with GATK (36) indel realignment. Long repeat sequences were annotated in the reference genome using LTRHARVEST (37) and REPEATMASKER (A.F.A. Smit, R. Hubley & P. Green RepeatMasker at http://repeatmasker.org) queried with repeat families identified by the GSC as well as a representative full length MAGGY LTR retro transposon. Large deletions were identified as regions of at least 300 bp with fewer than 10 reads per base pair interrupted by higher coverage regions of less than 50 bp. Deletions overlapping repeat annotations were removed, and the remaining per-strain deletions were normalized by clustering on genome location to generate a consistent set of deletion coordinates across strains. To identify short sequence variations, the full set of aligned reads was run through SAMTOOLS MPILEUP (http://www.htslib.org/doc/samtools-mpileup.html) and BCFTOOLS CALL (http://www.htslib.org/doc/bcftools.html). A variant allele was assigned to a strain if it was supported by at least 85% of the reads at the variant site for sites with at least 10 aligned reads for the given strain; variant sites where no allele passed these support criteria were flagged as ambiguous. Variant sites with zero or one alleles unambiguously assigned across all strains (*e.g.*, errors in the reference sequence) were removed as were sites overlapping annotated repeat regions or large deletions. Large deletions and variant sites were then classified among strains as summarized in Fig. 5.

### Southern hybridization of *vea1Δ* mutants

A 1052-bp probe to the 5’ UTR of *VEA1* was amplified from G217B *ura5^-^* genomic DNA using primer set OAS6659/OAS6660. A second 1013-bp probe to the *VEA1* CDS was amplified from G217B ura5^-^ genomic DNA using primer set OAS6663/OAS1969. In separate reactions the resulting PCR fragments were coupled with an alkaline phosphatase enzyme using the Amersham Gene Images AlkPhos Direct Labelling and Detection System (GE Healthcare), according to the manufacturers protocol. Genomic DNA from the parental strain G217B *ura5^-^*and the mutant strains was isolated by phenol chloroform extraction. In duplicate, 15 µg of genomic DNA from both samples was digested with restriction enzyme *Bgl*II for 1 hour at 37°C. Digested DNA was size separated by gel electrophoresis on a 0.7 % agarose gel and transferred to a Hybond-N+ nylon membrane (Amersham Biosciences). Membrane bound digested genomic DNA was hybridized at 60°C overnight with either the probe to the 5’ UTR or to the *VEA1* CDS. Finally, the membranes were visualized by using CDP- Star Chemiflourescent detection system (GE Healthcare).

### Flow cytometry

Transformants were selected at random and grown in liquid HMM with plasmid selection. Cultures were passaged 24 hours prior to harvest to obtain mid-log phase cells with an OD_600_ between 5-7. Approximately 1.25 E7 cells per sample were sonicated, washed, and spun down with D-PBS before being fixed on ice with BD Stabilizing Fixative (BD Biosciences). Once fixed, samples were washed and resuspended in D-PBS and analyzed using a BD LSRII Flow Cytometer in the UCSF Parnassus Flow Core. mCherry fluorescence signal in each sample was analyzed and quantified using FlowJo v. 10. mCherry positive and negative control strains were used to set gates for quantifying signal across all transformants.

## Acknowledgements

We thank Gregory Kujoth and Bruce Klein for sharing critical CRISPR reagents. We thank the UCSF Parnassus Flow CoLab RRID:SCR_018206 and DRC Center Grant NIH P30 DK063720 for flow cytometry instruments and support.

This work was supported by NIAID grant 2R37AI066224 to AS as well as funds from Chan Zuckerberg Biohub – San Francisco to AS. AS is a Chan Zuckerberg Biohub – San Francisco Investigator. The funders had no role in study design, data collection and interpretation, or the decision to submit the work for publication.

